# Investigating Brain and Biological Development in Children and Their Relationship with Physical, Mental, and Academic Outcomes

**DOI:** 10.1101/2024.10.07.617114

**Authors:** Hansoo Chang, Kevin Street, Ana Ferariu, Alexei Taylor, John Kounios, Fengqing Zhang

**Author notes:** Corresponding author: Fengqing Zhang Department of Psychological and Brain Sciences at Drexel University 3201 Chestnut St, Philadelphia, PA 19104.

## Abstract

Brain age and biological age, estimated using machine learning models with brain imaging and biological features, have emerged as promising biomarkers for predicting a broad range of health outcomes in adults. However, very few studies have examined the counterpart of brain age and biological age in children, that is Brain Development Index (BRDI) and Biological Development Index (BIDI). Existing studies on BRDI and BIDI are largely cross-sectional and do not provide adequate information on their temporal trajectory and predictive power for future health outcomes in children. Additionally, the interconnectedness of BRDI and BIDI across multiple health domains, especially child-specific developmental outcomes, remains underexplored. Our study utilized brain imaging features and blood-based biomarkers from the Adolescent Brain and Child Development (ABCD) study to assess the trajectory of BRDI and BIDI over multiple time points. We examined their relationships with physical, mental, and academic health outcomes.

Lastly, we utilize Bayesian network analysis to examine the relationship between the two indexes and their subcomponents. We found that delayed BRDI and BIDI were significantly associated with adverse future health outcomes across several domains. In addition, Bayesian network analysis revealed BRDI and BIDI subcomponents influence one another across different organ systems. Additionally, males exhibited more advanced BRDI, while females showed more advanced BIDI, revealing important sex differences in adolescent development. This research provides the first comprehensive analysis of BRDI and BIDI trajectories, revealing their predictive power for future health outcomes and offering new insights into the interconnected development of brain and biological systems in children.

## 1. Introduction

Whole-person health reflects the complex interplay of brain, biological, lifestyle, and social factors affecting an individual’s well-being (Tilson et al., 2020). Likewise, brain age and biological age, estimated using machine learning models with brain imaging and biological features respectively, have emerged as promising biomarkers for quantifying whole-person health and predicting a broad range of health outcomes beyond chronological age in adults (Taylor et al., 2022; Zhang et al., 2023). Individuals of the same chronological age can differ in their brain or biological age, indicating varying levels of risk for health conditions (Liang et al., 2019; Niu et al., 2022; Niu et al., 2020). However, very few studies have examined the counterpart of brain age and biological age in children, that is Brain Development Index (BRDI) and Biological Development Index (BIDI) (Tian et al., 2023).

Similar to brain age, BRDI is commonly estimated using brain imaging features (e.g., gray matter volume) since physiological differences in these brain features can affect social and physical outcomes in adolescents (Franke & Gaser, 2019; Richards & Xie, 2015). Physiological changes of grey matter volume (GMV) during early development are associated with cognitive delay (Erus et al., 2014), ADHD (Giedd et al., 2015), childhood-onset schizophrenia (Giedd et al., 2015), poverty (Hair et al., 2015), and academic achievement (Hair et al., 2015) showing that BRDI may be useful tool in predicting these child-related health outcomes. Similarly, BIDI is often estimated using machine learning models and blood-based and anthropometric biomarkers (e.g., blood pressure (Flynn et al., 2017), BMI (Geserick et al., 2018)) that are indicative of different organ development in the body to predict chronological age. Existing studies have shown that BIDI is associated with a variety of physical health outcomes such as childhood hypertension (Laurent et al., 2019), early-life adversity (Colich et al., 2020), cardiorespiratory problems, physical problems, and fitness (Sitovskyi et al., 2019).

Although these biomarkers are promising, our study is motivated by the significant gaps remaining in the understanding of BRDI’s and BIDI’s role in childhood development over time. Existing studies are largely cross-sectional and do not provide sufficient evidence on BRDI’s ability to predict future health outcomes in children (Niu et al., 2020). Few studies have examined BRDI longitudinally in adolescents, making it necessary to investigate the normal trajectory of brain development over time (Tian et al., 2023; Zhao et al., 2022). While large datasets, like the UK Biobank, have been used to study brain aging in adults (Cole, 2020; Tian et al., 2023), similar research in children remains limited. Additionally, the relationship between BRDI and physical or child-specific outcomes, such as school grades, has not been thoroughly explored (Cole et al., 2019; Liem et al., 2017), and there is no normative BRDI growth chart for children, leaving the typical developmental trajectory unclear (Bethlehem et al., 2022).

Similarly, research on the Biological Development Index (BIDI) is scarce, particularly in relation to child-relevant outcomes (Colich et al., 2020). While biological age in adults has been linked to neurological and social outcomes (Belsky et al., 2015), it is unknown whether BIDI can predict broader health outcomes in children. Furthermore, to our knowledge, no effective BIDI growth chart exists, making it challenging to assess children’s physical development across different ages (Sanders et al., 2015). Addressing these gaps requires longitudinal data to explore the trajectories of BRDI and BIDI and their predictive value for health outcomes in children.

Lastly, existing studies often treat these indexes separately, overlooking the interconnectedness of body systems especially as recent papers have shown proof of one affecting the other (Feldman & Eidelman, 2009; Tian et al., 2023; Turesky et al., 2019). In adults, greater biological age has been associated with lower total brain volume, linking multi-organ decline to nervous system health (Whitman et al., 2024), but whether these relationships exist in children remains unknown. Furthermore, it is unclear how BRDI and BIDI influence each other in childhood development. A dual approach is critical, to understand if changes in brain structure can affect or be affected by changes in biological markers. For example, UK Biobank data showed that brain and biological age influence each other’s rate of acceleration over time in adults (Tian et al., 2023). A combined analysis could provide essential insights into the mechanisms of development and whole-person health in children.

In this current study, we utilized brain imaging and blood-based biomarkers across multiple time points to calculate the acceleration and trajectory of BRDI and BIDI in a cohort of children across the United States in the Adolescent Brain and Child Development (ABCD) study, the largest long-term study of brain development in child health in the United States. In addition, we comprehensively examined the relationship between BRDI and BIDI with a multitude of physical, mental, and academic health outcomes. Lastly, we utilize Bayesian network analysis to examine the relationship between the two indexes and their subcomponents. Our study elucidates the trajectory of normal brain and biological development in children over several years and reveals the predictive ability of BRDI and BIDI across several health domains which to our knowledge, no study has yet done. These analyses in addition to our Bayesian network analysis that finds probabilistic relationships between BRDI, BIDI, and their subcomponents, hold great potential to improve our understanding of normal child brain and biological development and how abnormal development may affect future health outcomes that may otherwise have been unknown and unquantifiable.

## 2. Materials and Methods

### 2.1 Participants

We used data from the 5.1 release of the Adolescent Brain Cognitive Development (ABCD) study, the largest longitudinal study of brain development and youth health in the United States. Data was available at three time points (baseline, 2-year, and 4-year) for the BRDI calculations and at two time points (2-year and 4-year) for quantifying BIDI. Most outcome health variables had data available up to the 4^th^ year of data collection unless otherwise stated.

The baseline ABCD cohort was represented by 11,868 participants (52% white, 52% males) coming from 9807 unique families across 22 sites with mean age of 119 months (9-10 years old, SD = 7.5). Any participants who had a preterm birth were removed. Tables S3 and S4 in the appendix details the age, race, and sex distribution at each time point.

### 2.2 Measures

#### 2.2.1 Brain Development Index (BRDI)

To calculate BDI, structural MRI gray matter volumes using the Desikan atlas from the ABCD 5.1 Data Release were utilized (Desikan et al., 2006). Structural MRI GMV is commonly used in BRDI calculations (Cole et al., 2018), since gray matter integrity has been shown to change at the microscopic and macroscopic level during child development (Wang et al., 2019). In addition, GMV rather than gray matter thickness was utilized since GMV alone provided more accurate estimates of BRDI than gray matter thickness or gray matter thickness and volume together in our analyses. Procedures for image acquisition, image processing, and quality assurance have been described previously in the ABCD documentation (Casey et al., 2018). The present analyses included 68 cortical regions that were derived from the surface-based atlas procedure developed by Desikan et al. (2006) in addition to 19 subcortical regions that were derived by the automated labeling procedure developed by Fischl et al. (2002).

In accordance with the literature, stepwise AIC feature selection was applied to exclude regions of interest (ROIs) not correlated with chronological age, reducing noise in the BRDI calculations (Franke et al., 2012). Feature selection focused on healthy participants (N = 1325) to identify ROIs reflecting typical age-related development, with the final ROI list detailed in Table S1. BRDI was calculated using the Klemera-Doubal Method (KDM), a robust method for assessing deviations of biomarkers from age-related norms (Klemera & Doubal, 2006), suitable for this study’s narrow age range at baseline. The trained model was validated on a healthy test set (N = 1322) using Root Mean Squared Error (RMSE) and Mean Absolute Error (MAE). It was then applied to unhealthy participants and follow-up data to calculate BRDI and BRDI Gap, indicating advanced or delayed brain development. Further details on the BRDI pipeline and criteria are available in the supplementary materials.

#### 2.2.2 Biological Development Index (BIDI)

Participants with available biomarkers necessary to calculate biological development index were also split into healthy and unhealthy based on the same criteria used for brain development index. For similar reasons as BRDI, we chose KDM to calculate BIDI. Additionally, KDM has been shown to be a promising and accurate method for estimating biological age, which has been utilized by prior key biological aging papers (Belsky et al., 2015). Procedures for physical health data collection are described previously (Uban et al., 2018). Blood samples are taken during the 2-year and 4-year follow-up waves, with no changes in methodology since the start. A full list of blood biomarkers used for calculating BIDI are shown in Table S2 in the supplement.

BIDI was also calculated using KDM and trained on the healthy subset of participants at the 2-year follow up (N = 210) since the blood-based biomarkers were not collected at baseline. The trained model was projected to the healthy test set at the 2-year follow up to assess model accuracy using RMSE and MAE. Then, the trained model was used to project the BIDI values to the unhealthy 2-year follow up participants, and all the 4-year follow up participants.

Additionally, we computed the difference between the predicted BIDI and chronological age, called BIDI Gap, to capture advanced or delayed biological development. A healthy participant is expected to have a BRDI and BIDI Gap close to zero since healthy participants’ predicted development index should be on average the same as their chronological age in “normal” development.

#### 2.2.3 Physical Health Outcomes

The ABCD data also administered a medical history questionnaire annually since baseline to assess if participants have developed any medical issues ranging from asthma to cancer. These outcomes were coded as “1” if the participant has had the health condition since the last follow up time point (every two years), and a “0” if the participant has not had the health condition since the last time point.

#### 2.2.4 Mental Health Outcomes

A wide range of mental health outcomes were used as outcomes to assess the association between BRDI and BIDI and child mental health outcomes. Briefly, we used the parent-report KSADS, which is a semi-structured interview to measure current and past symptoms of mood, anxiety, psychotic, and disruptive behavior disorders, among others, in children ages 6-18 years old (Townsend et al., 2020). Furthermore, the 7-Up Mania Inventory was utilized to assess youth mania symptoms (Youngstrom et al., 2013), and the CBCL, an ASEBA instrument assessing youth dimensional psychopathological syndromes dimensional and adaptive functioning, was used to examine broad psychopathology (Achenbach, 2009). Lastly, alcohol sips and alcohol latent classes were analyzed to examine the relationship between the indexes and alcohol usage. To assess early alcohol exposure, we used data measured by the iSay Sip Inventory, including questions regarding age of alcohol use onset, number of sips and type of alcohol ingested (Jackson et al., 2015). The alcohol latent classes were created using previous work from Ferariu et al. (2024) which display distinct personality traits (Ferariu et al., 2024).

#### 2.2.5 Academic Outcomes

Lastly, we analyzed academic outcomes at 3-year follow up to understand the relationship between the development indexes and a health outcome that is specific to children rather than adults. We used the School Attendance and Grades information in the ABCD data which provided information on how often the participants would skip class and their most recent school grades (Zucker et al., 2018). Parents and students were asked how many total days the participant skipped school without an excuse. Grade estimates as provided by the ABCD study range between 1 (corresponding to A+) and 12 (corresponding to a failing grade) (Zucker et al., 2018).

### 2.2 Statistical Analyses

Mixed-effects models using the lme4 package in R were utilized to examine whether BRDI Gap at baseline or BIDI Gap at 2-year follow up prospectively predicted the health outcome of interest at the latest available time point (either 4-year follow up or 2-year follow up) (Bates, 2010). Data from the training sets used to estimate BRDI and BIDI were excluded from this set of analyses to avoid information leakage and overfitting. For non-normally distributed outcomes, generalized mixed-effects models were used (e.g., mixed-effects logistic regression was used for binary outcomes and a zero-inflated mixed-effects model using the cplm package in R was used for the outcomes with a zero-inflated distribution) (Jackman, 2010). The interaction between the development index gap and sex was also included to account for the biological differences between sexes and test whether the effect of BRDI and BIDI on a health outcome was moderated by sex. Furthermore, the outcome measure at baseline, sex, race, and SES were included as covariates. The random effects for models with BIDI included family ID, site ID, while the random effects for BRDI models includes family ID, site ID, and MRI manufacturer. These random effects were included to account for participants nested within families and geographic locations.

To examine heterogeneous development across multiple organ systems, we conducted Bayesian network analysis using the bnlearn package in R to examine the relationships between BRDI and BIDI across different organ systems (Scutari, 2009). Following literature recommendations (Tian et al., 2023), we calculated BIDI for different physiological systems including cardiovascular, organ function, musculoskeletal, immune, and metabolic health. For each physiological category in the BIDI, we calculated an ’organ development index’ using KDM to represent that system’s maturity. Similarly, we calculated multiple brain development indexes using different grey matter measurements (i.e., grey matter thickness, and both grey matter volume and thickness). We utilized blood-based sample and structural MRI data at the 2-year follow-up as this was the first timepoint in which we could calculate both BRDI and BIDI. Using Bayesian network analysis, we examined the complex relationships between BIDI, multiple organ development indexes, and BRDI to explore how biological development is interconnected with and can influence and be influences by brain development trajectories.

## 3. Results

### 3.1 Brain Development Index and Biological Development Index in Adolescents

#### 3.1.1 Prediction of Brain Development Index

The predicted BRDI ranged from 5.21 to 15.83 years, with a mean of 10.03 and a standard deviation of 1.46 years at baseline (chronological age mean = 10.10, SD = 1.60). For the healthy test set, the mean was 9.87 (SD = 1.02), while the unhealthy set had a mean of 9.88 (SD = 1.11). The BRDI distribution was normal, and the mean BRDI Gap (difference between BRDI and chronological age) was –0.05 (SD = 0.83 years).

The model accurately predicted BRDI for healthy participants at baseline, achieving an RMSE of 0.52 years and MAE of 0.39 years in the training set. For unseen healthy participants, the RMSE was 1.03 years, and MAE was 0.83 years; for the unhealthy group, RMSE was 1.39 years and MAE was 1.10 years. Notably, our model outperformed studies like Cole et al. (2018) and Holm et al. (2023), which had higher MAEs. The strong performance is likely due to training on participants with a narrow age range (9-10 years). For the 2-year follow-up, RMSE was 1.43 years and MAE 1.14 years, while at 4 years, they were 1.60 and 1.30 years, respectively.

Partial R-squared analysis identified key regions for BRDI calculations, including the superior parietal lobule, ventral diencephalon, and left amygdala. A greater GMV in the superior parietal lobes and left inferior parietal lobe reduced BRDI estimates, while larger GMV in the diencephalons and left amygdala increased BRDI predictions. A full list of feature importance is provided in Table S1. (Holm et al., 2023)

#### 3.1.2 Prediction of Biological Development Index

The predicted BIDI ranged from 8.76 to 14.20 years, with a mean of 11.89 years and a standard deviation of 0.80 years for all participants excluding the training set (chronological age mean = 11.92, SD = 0.66) at the 2-year follow-up. The healthy test set had a mean BIDI of 11.95 (SD = 0.80), while the unhealthy set had a mean of 11.81 (SD = 0.80). The BIDI distribution was normal, and the mean BIDI Gap (difference between BIDI and chronological age) was –0.06 (SD = 0.65 years).

The model accurately predicted BIDI for healthy participants at the 2-year follow-up, achieving an RMSE of 0.52 years and MAE of 0.39 years in the training set. For unseen healthy participants, the RMSE was 0.64 years, and MAE was 0.49 years; for the unhealthy group, RMSE was 0.68 years and MAE was 0.51 years. This performance is notable compared to a reference study by Putin et al. (2016), which achieved an MAE of 6.07 years using similar biomarkers. At the 4-year follow-up, the model’s RMSE was 1.11 years, and MAE was 0.85 years.

Partial R-squared analysis identified key blood-based biomarkers for BIDI, including hemoglobin, mean arterial pressure, platelet count, and BMI. A hemoglobin, mean arterial pressure, and BMI increased BIDI estimates, while a higher platelet count decreased BIDI. Further details are provided in Figure 1.

**Figure 1.**
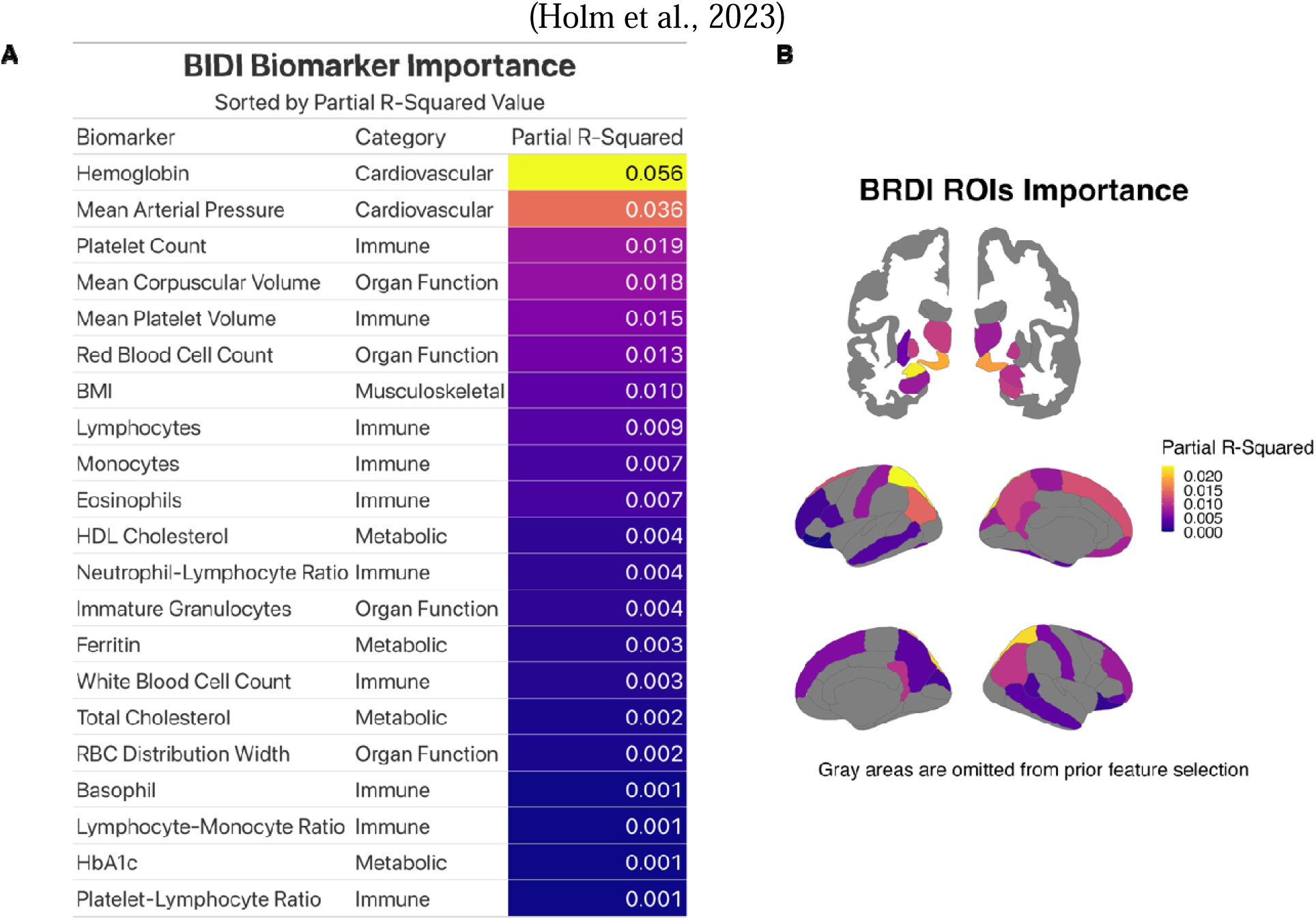
Individual biomarker importance in BRDI and BIDI. A: BIDI biomarkers ordered by partial R-squared for BIDI prediction for all participants. Hemoglobin and mean arterial pressure were the two most important predictors in our BIDI model as they both had a medium effect size. B: GMV ROIs ordered by partial R-squared for BRDI prediction for healthy participants. Gray areas are ROIs omitted by Step AIC due to low correlation with chronological age. Left hemisphere subcortical ROIs and the superior parietal lobe in both hemispheres were the most important ROIs in calculating BRDI.

#### 3.1.3 Distribution of BRDI and BIDI

To understand the trajectory of brain and biological development, we developed BRDI and BIDI growth charts similar to BMI growth charts used by clinicians. As shown in Figure 2, BRDI increased steadily over the years for healthy participants. However, males on average had a more advanced BRDI than females at the same chronological age. This finding is also consistent in the brain age literature where men are found to have older brain ages compared to females.

**Figure 2.**
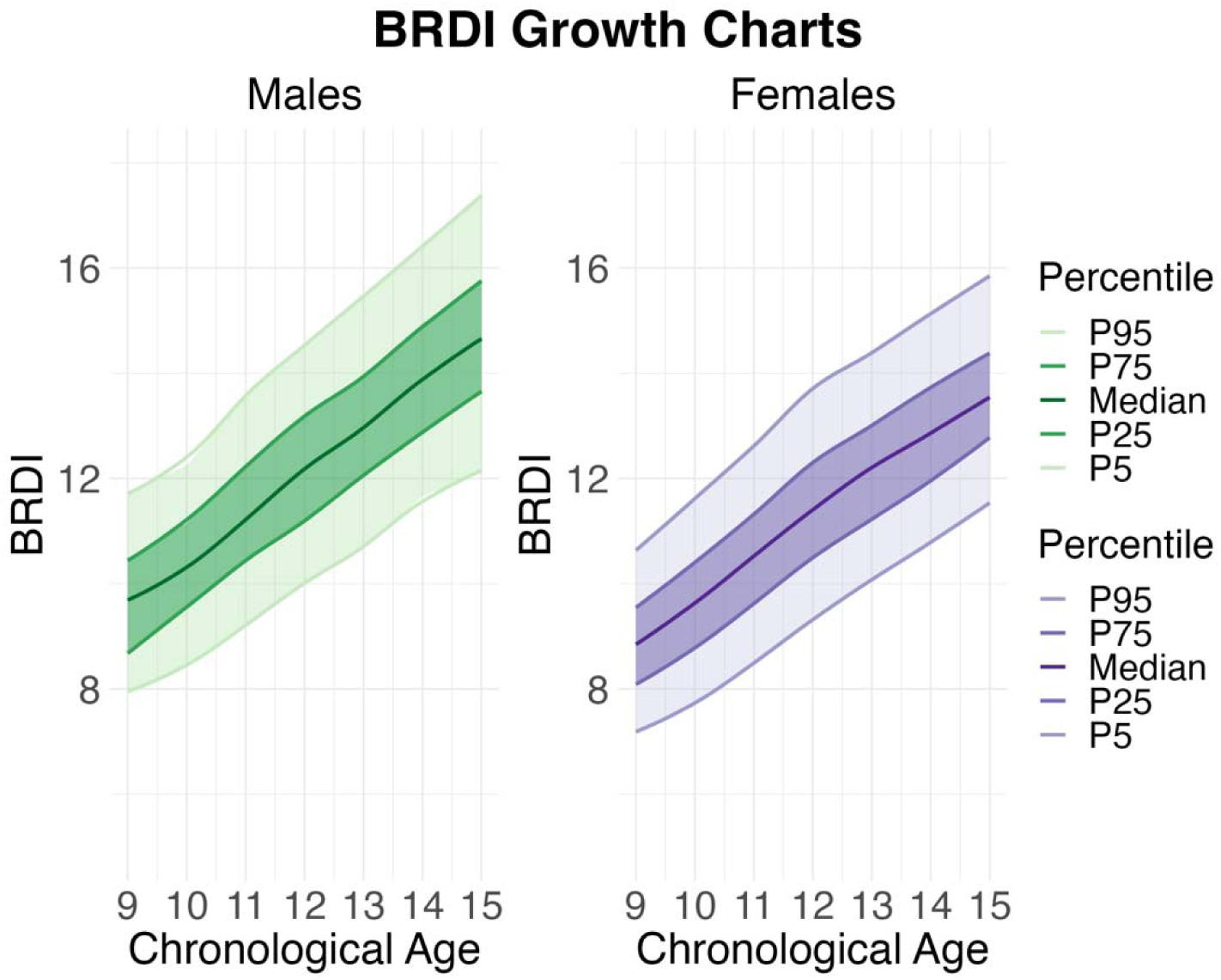
BRDI Growth charts for healthy males and females. Males on average had a more advanced BRDI relative to girls of the same age. BRDI for both sexes followed a linear path, in which BRDI roughly equals the chronological age at a given year although within a chronological year. There is about a 4-year BRDI range between the 95^th^ and 5^th^ percentile.

We also developed the BIDI growth chart as shown in Figure 3. BIDI increased steadily over the years for healthy participants. However, the ranges for the percentiles may not be as stable as the BRDI Growth Chart due to a smaller sample size. Females on average had a more advanced BIDI than males at the same chronological age. In addition, BIDI had only two time points relative to BRDI which has 3 time points, resulting in a smaller range for chronological range in the chart.

**Figure 3.**
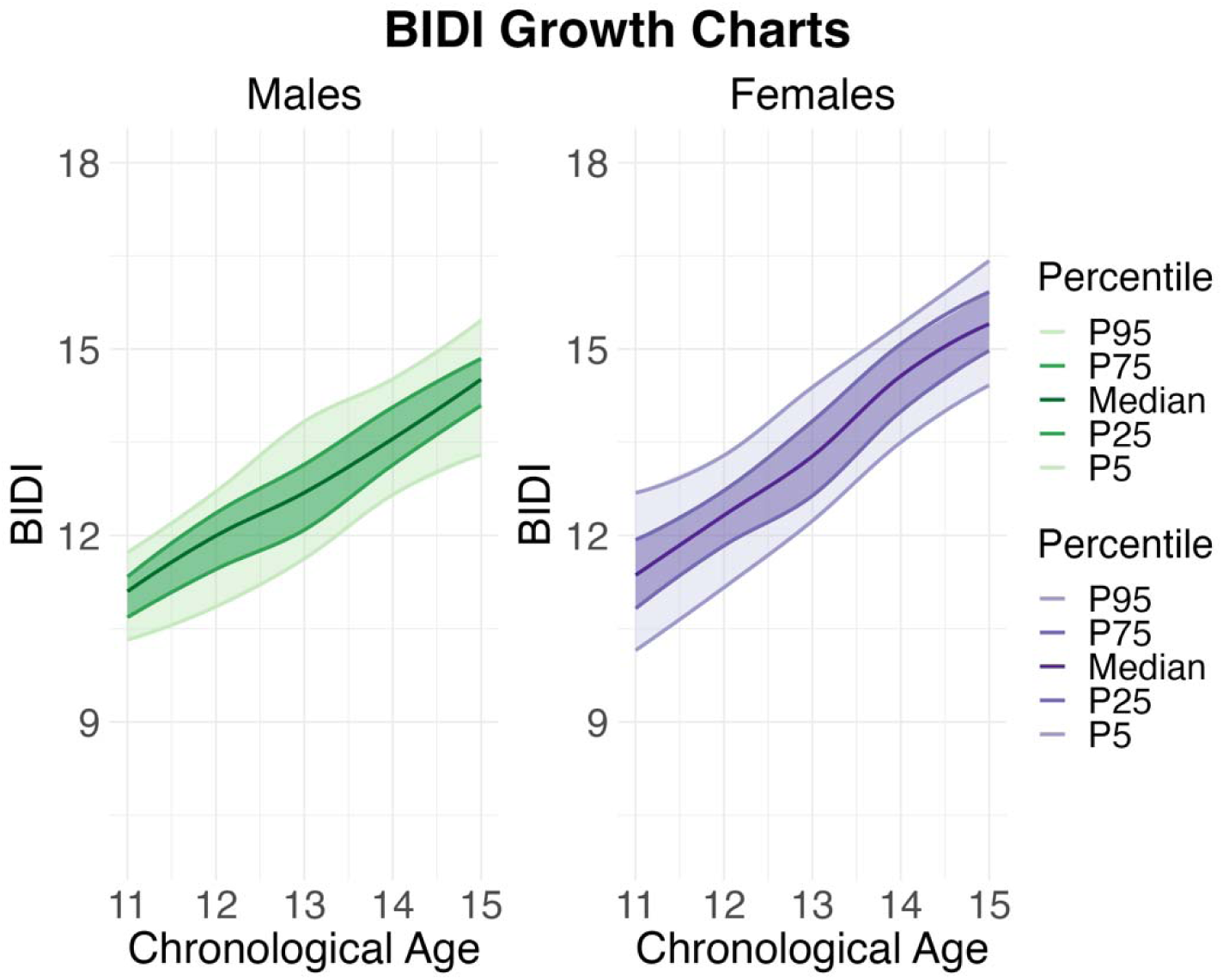
BIDI Growth Chart for healthy males and females. Females on average had a more advanced BIDI relative to males of the same age. BIDI for both males and females follow a linear path in which BIDI roughly equals the chronological age at a given year although within a chronological year. There is a smaller range between the 5^th^ and 95^th^ percentile compared to the range of BIDI.

To illustrate the potential clinical utility of these growth charts, we explored whether individuals above the 95^th^ percentile or below the 5^th^ percentile of BRDI showed distinct health profiles. As shown in Figure 4, participants with an advanced BRDI, that is, being above the 95th percentile, had less health problems across three health domains particularly the neurocognitive domain. Furthermore, having a delayed BRDI, or being under the 5th percentile was associated with more health problems particularly in the neurocognitive domain. Specifically, those below the 5th percentile in BRDI at baseline had significantly more total KSADS diagnoses two years later relative to those within the 95^th^ and 5^th^ percentile (Figure 4). Detailed results on the association between BRDI or BIDI and a broad range of child health outcomes are summarized in the following section.

**Figure 4.**
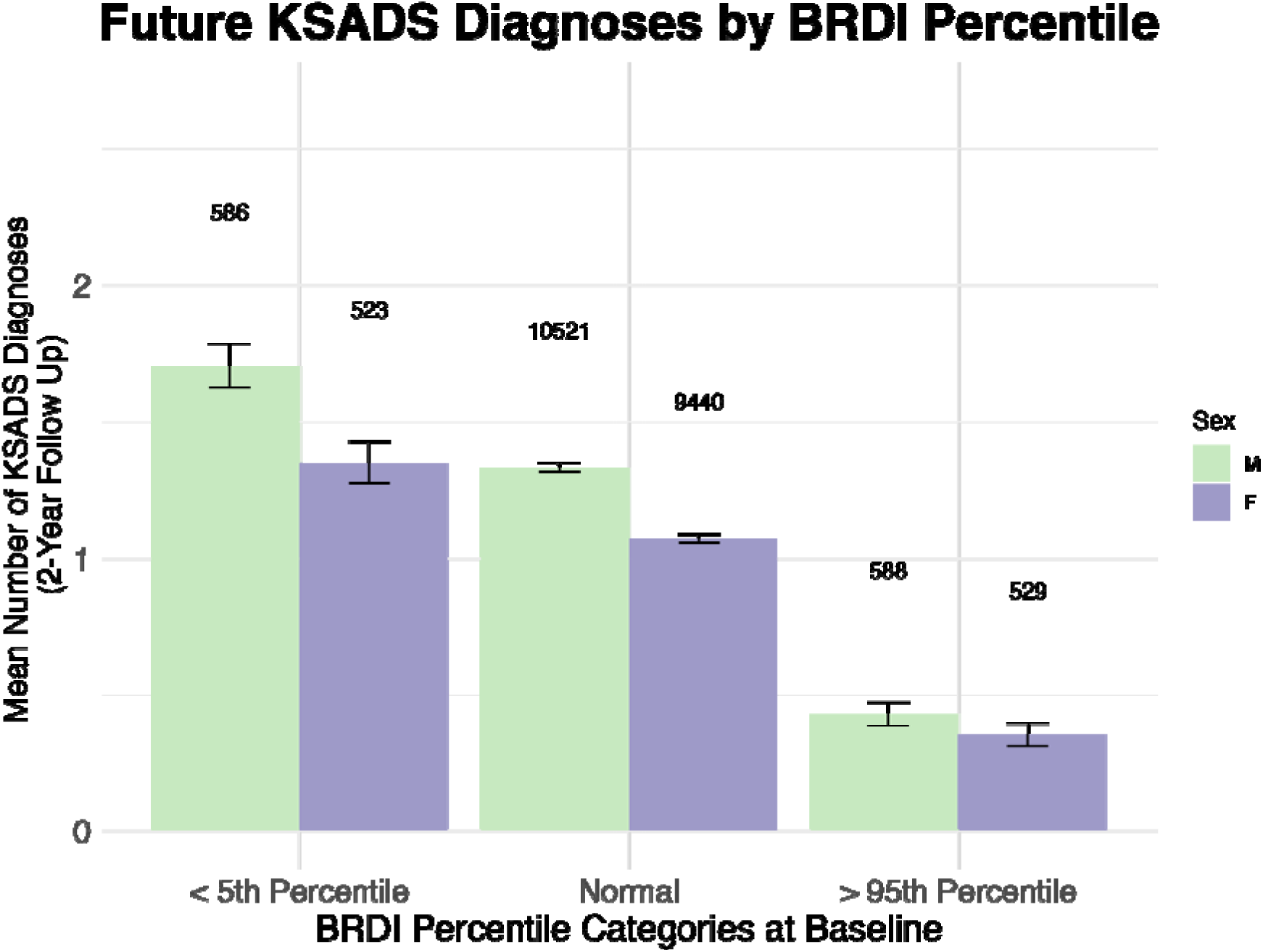
Delayed BRDI (< 5^th^ Percentile) at baseline for both males and females were more likely to have more KSADS mental health diagnoses 2 years later than those within the 5^th^ and 95^th^ percentile. In addition, those with an advanced BRDI (> 95^th^ Percentile) were on average found to have less KSADS diagnoses two years later than both those with a delayed BRDI and those within the 95^th^ and 5^th^ percentile range.

### 3.2 BRDI and BIDI Predicting Health Outcomes

Across the physical, mental health, and academic outcomes, we found that delayed BRDI was significantly associated with worse health outcomes at future time points. Alcohol latent class and KSADS OCD were significantly associated with advanced BRDI on the other hand.

Likewise, delayed BIDI was associated with worse health outcomes across the three different health domains. Delayed BRDI and BIDI were both predictive of worse school grades. In general, BIDI was predictive of different health outcomes than BRDI such as respiratory, eye and ear problems, and CBCL depression. These results are illustrated in Figure 5 with detailed descriptions in the following subsections.

**Figure 5.**
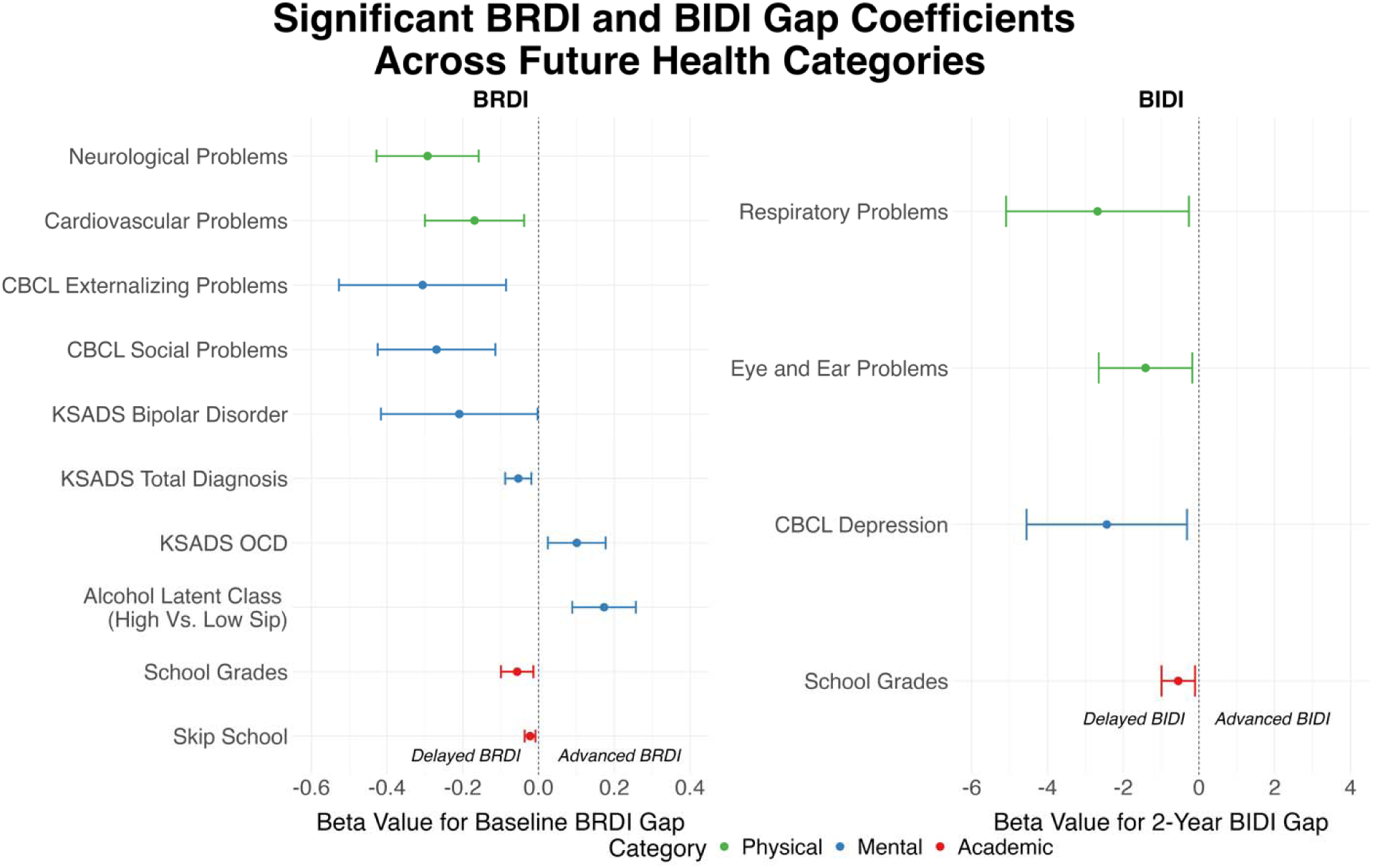
Significant BRDI and BIDI Gap regression coefficients for future health outcomes (either 2-year or 4-year follow up). Beta coefficients were standardized to make the regressions outputs comparable. All health outcomes were scaled such that a higher value is associated with a worse health outcome. Overall, delayed BRDI and BIDI were associated with worse health outcomes in the future. Both indexes were associated with different health outcomes in the three different health domains.

#### 3.2.1 Physical Outcomes

##### Brain Development Index

For physical health outcomes at the latest possible time point (4-year follow up), baseline BRDI Gap was a significant predictor of cardiovascular problems (b = -0.169, *p* = .011) and neurological problems (b = -0.293, *p* < .005) at the 4-year follow up. A delayed BRDI was associated with a higher risk of developing cardiovascular and neurological problems 4 years later as the beta coefficients for both significant outcomes are negative.

##### Biological Development Index

BIDI Gap at 2-year follow up was used to examine 4-year follow up health outcomes since 2-year follow up BIDI Gap was the earliest available time point. BIDI Gap was a significant predictor of eye and ear problems (b = -1.411, *p* = .032) and respiratory problems (b =-2.678, *p* = .030) at the 4-year follow up. A delayed BIDI was associated with a higher risk of developing eye and ear and respiratory problems 2 years later as the beta coefficients for both significant outcomes are negative.

#### 3.2.2 Mental Outcomes

##### Brain Development Index

All significant BRDI Gap beta coefficients were negative for the neurocognitive outcomes, suggesting that a delayed BRDI at baseline was significantly associated with neurocognitive outcomes at the latest time point (2-year follow up). Since the latest KSADS data was available at the 2-year follow up, baseline BRDI Gap was used to predict these two-year scores. Overall, BRDI Gap had a significant beta value for the overall total KSADS summary score (b = -0.053, *p* = .003). Of the available measures in the KSADS at the 2-year follow up, the effect of BRDI Gap on the likelihood of generalized bipolar disorder at 2-year follow-up was significant (b = -0.209, *p* = .047). Furthermore, the effect of BRDI Gap on the likelihood of obsessive-compulsive disorder (OCD) at 2-year follow-up was significant (b = 0.101, *p* = .009). Specifically, we observed that a delayed BRDI at baseline was significantly associated with a higher likelihood of developing generalized bipolar disorder and having more KSADS problems in the future whereas an advanced BRDI was associated with worse OCD outcomes in the future.

In addition, we found that baseline BRDI Gap was a significant predictor of CBCL social problems (b = -0.269, *p* = .032) at 4-year follow up and externalizing problems (b = -0. 306, *p* = 0.039) at 4-year follow up, which is indicative of rule-breaking and aggression. Since the latest time point for CBCL outcomes is the 4-year follow up making these outcomes a 4-year prediction.

Briefly, alcohol latent classes were categorized into “No sip,” “Low sip,” and “High sip” based upon work done by Ferariu et al. (2024). Baseline BRDI Gap was found to be predictive of differentiating between latent class 3 (High sip) and latent class 2 (low sip) from baseline to 4- year follow up. BRDI Gap at baseline was a significant predictor of being within the “high sip” latent group relative to the “low sip” latent group, with no significant differences between sexes in this relationship. An advanced BRDI was associated with a higher likelihood of being in the high sip group relative to the low sip group.

Lastly, although we did not observe a significant beta coefficient for baseline BRDI Gap on the 4-year score of summed score of Mania of the 7-up Mania Inventory, we observed a significant interaction effect between baseline BRDI Gap and being male (b = -0.132, *p* = .013) indicating that the interaction between BRDI Gap and mania is moderated by sex. Specifically, it would suggest that the effect of having an advanced BRDI is diminished in males compared to females when it comes to mania problems in the future.

##### Biological Development Index

BIDI Gap at 2-year follow up was a significant predictor of CBCL Depression on the DSM-V scale (b = -2.434, *p* = .026). Specifically, a delayed BIDI is associated with a greater likelihood of developing depression at future timepoints. BIDI Gap was not a significant predictor of the 7-up mania sum score, and the KSADS measurements latest available time point was at the 2-year follow up time point, meaning that the BIDI analysis at 2-year would not be a predictive analysis.

#### 3.2.3 Academic Outcomes

##### Brain Development Index

2-year BIDI Gap was found to be a significant predictor of school grades at the 4-year follow up (b = -0.547, *p* < .001). We observed that a delayed BRDI was associated with worse overall school grades 2 years into the future. In addition, participants with a delayed BRDI at baseline were more likely to skip school at the 3-year follow up (b = -0.022, *p* = .002).

##### Biological Development Index

2-year BIDI Gap was found to be a significant predictor of school grades at the 4-year follow up (b = -0.547, *p* < .001). We observed that a delayed BIDI was associated with worse overall school grades 2 years into the future.

### 3.3 Bayesian Network Analysis

Results from our Bayesian network analysis demonstrated the connectedness of BRDI to BIDI through the subcomponents and vice versa at 2-year follow up (Figure 6). After running 10- fold cross validation optimizing for log-likelihood loss, Incremental Association Markov Blanket (IAMB) was found to be the most optimal constraint-based algorithm. The learned Bayesian network included 9 nodes and 18 directed arcs, with an average Markov blanket size of 4.89 and an average neighborhood size of 4.00. The branching factor was 2.00, indicating that each index affects two other indexes on average. Only arcs with a significant p-value (*p* < 0.05) are drawn in Figure 6, and BIC was used to determine arc strength. Organ function development index influenced both BRDI and BIDI directly in addition to other subcomponents revealing an intricate system of whole-person health. Therefore, although BRDI and BIDI may not affect each other directly, these two indexes may be influenced by their subcomponents (organ function) and likewise influence each other via different subcomponents (ex. cardiovascular index).

**Figure 6.**
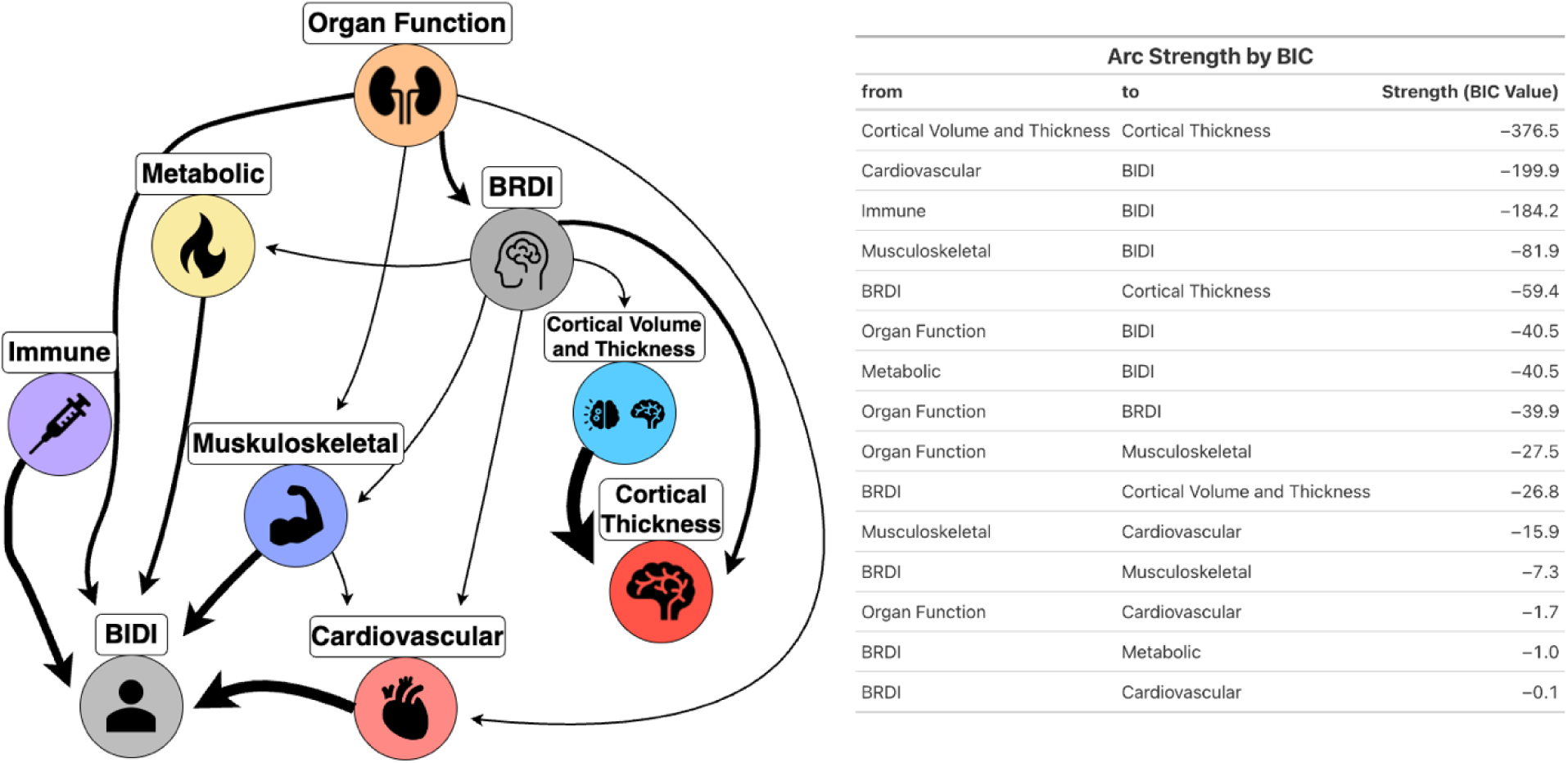
Bayesian Network Analysis Directed Acyclic Graph (DAC) of BRDI, BIDI, and their sub-indexes. Significant BIC values are shown. The relationships between BRDI and BIDI are more clearly defined through their subcomponents. As an example, organ function index influences BRDI and BIDI directly while BRDI influences BIDI indirectly through the musculoskeletal index.

## 4. Discussion

Our research has developed a tool using grey matter volume and blood-based biomarkers to predict adolescent health outcomes across physical, mental, and academic domains. We created the first BRDI and BIDI growth charts for adolescents, akin to BMI growth charts used by clinicians. These charts track BRDI and BIDI development which has not been done before and were shown to hold potential in predicting mental health outcomes such as KSADS diagnoses. A delayed BRDI was found to be predictive of adverse future health outcomes which were supported by our multilevel regression analyses, demonstrating the indexes importance and consistency in assessing whole-person health. Our Bayesian network analysis also revealed the interconnectedness of BRDI, BIDI, and their subcomponents suggesting mutual influence. Our study advances the understanding of adolescent brain and biological development and provides clinicians with a practical tool to predict future health outcomes, enhancing adolescent healthcare.

Our models outperformed previous studies like Cole et al. (2018), with validation metrics that underscore their robustness. For example, growth charts showed that males often have a more advanced BRDI but delayed BIDI compared to females, consistent with previous findings (Belsky et al., 2015). A narrower BIDI range suggests more stable blood biomarkers over time (Nguyen et al., 2019). Cardiovascular markers like arterial pressure were important for BIDI (Theodore et al., 2015), while HbA1C showed limited utility due to its stability in childhood (Lind et al., 2019) highlighting that certain biomarkers may be more important for BIDI in children than in adults since HbA1C is generally an important biomarker in biological aging in older adults (Belsky et al., 2015).

Not only did our models perform better on statistical metrics than previous studies, but our BRDI and BIDI also successfully predicted child-relevant health outcomes. Overall, we found that delayed BRDI correlates with future neurological and cardiovascular issues, while delayed BIDI is linked to respiratory and eye and ear conditions. This aligns with adult studies on brain age and health (Cole et al., 2019; Tian et al., 2023), marking our work as one of the first to observe similar patterns in children. Higher eosinophil counts associated with delayed BIDI were consistent with respiratory conditions, like asthma, as highlighted by other research on adolescents (Shamriz et al., 2018; Trivedi & Denton, 2019). Additionally, delayed BRDI and BIDI were associated with poorer school performance, aligning with research linking smaller subcortical GMV to academic struggles (Hair et al., 2015; Urrila et al., 2017). Our study is among the first to explore academic outcomes using BRDI specifically. BIDI’s association with stress and lower socioeconomic status (SES) may further explain its impact on school performance (Heissel et al., 2017). These outcomes reveal the utility of BRDI and BIDI in predicting whole-person outcomes that align with previous findings the examine these outcomes individually.

The Bayesian network analysis highlighted a moderately connected structure, emphasizing links between BRDI, BIDI, and organ function, supporting findings from the UK Biobank (Mishra et al., 2023). Our findings showed that while BRDI and BIDI may not directly influence each other, their relationship is mediated through shared subcomponents like the organ function index, which directly impacts both BRDI and BIDI. For example, the organ function index connects to BRDI and BIDI, suggesting that changes in organ health can have cascading effects on both brain and biological age. Furthermore, BRDI was found to influence BIDI indirectly through the musculoskeletal index, highlighting a pathway through which brain development can impact broader physiological systems. This interconnected structure emphasizes that whole-person health is not simply a sum of its parts but rather an intricate network where multiple systems influence each other. However, our study’s short follow-up period and cross-sectional Bayesian analysis limit long-term insights. Future research should examine BRDI and BIDI through adolescence into adulthood, particularly through puberty. A dynamic analysis across multiple time points could further clarify the biological relationship between BRDI and BIDI over time.

In summary, our study provides a valuable tool for clinicians and researchers to monitor adolescent development. By introducing the first BRDI and BIDI growth charts and demonstrating their predictive value, we offer a new perspective on adolescent brain and biological health.

## Notes

### Competing Interest Statement

The authors have declared no competing interest.

