## Appendix for "Investigating Brain and Biological Development in Children and Their Relationship with Physical, Mental, and Academic Outcomes"

Measures

In accordance with the literature (Franke et al., 2012), we applied stepwise AIC for feature selection to exclude any regions of interest (ROIs) not correlated with chronological age, reducing noise in the BRDI calculations and enhancing the accuracy of the prediction. The finalized ROI list is detailed in Table S1 of the supplement. We segregated participants into healthy and unhealthy groups using baseline Child Behavior Checklist (CBCL ASEBA) and Medical History reports, which encompass mental and physical health.

The BRDI model was trained on the training set of the healthy participants at baseline (N = 1325). We anticipate that the healthy participants’ predicted BRDI will be closer to their chronological age relative to the unhealthy participants’ predicted BRDI. Feature selection was conducted solely on healthy participants to pinpoint ROIs that most reliable reflect typical age-related development. We utilized the Klemera-Doubal Method (KDM) to calculate our BRDI since this method has been shown to be a robust method for assessing the deviation of an individual's biomarkers from established age-related norms (Klemera & Doubal, 2006). We chose KDM because it does not regress the ROIs on chronological age like many other biological age estimation algorithms. Since participants in the ABCD study were all 9-10 years old at baseline allowing little variation in their chronological age, any algorithms regressing biomarkers on chronological age are unlikely to yield accurate estimation. Although KDM was first developed to calculate biological age, this method can also be applied to calculating brain age since the algorithm does not discriminate between biomarkers if its value changes with chronological age. Furthermore, researchers interested in aging indexes have commonly cited this method as a statistical method rather than strictly a biological aging method (Cole et al., 2019). Similar to blood-based biomarkers, gray matter volume may also be treated like a blood-based biomarker regarding calculating BRDI using KDM.

This calculated BRDI was then projected to a healthy test set at baseline (N = 1322) to test model accuracy using Root Mean Squared Error (RMSE) and Mean Absolute Error (MAE). After the model accuracy was verified, the trained model was used to project BRDI for the unhealthy subset at baseline, all participants at 2-year follow up, and 4-year follow up. Additionally, we computed the difference between the predicted BRDI and chronological age, called BRDI Gap. A positive or negative BRDI gap represents advanced or delayed brain development. Details of the BRDI pipeline and exclusion-inclusion criteria are explained in the supplementary materials.

Table S1

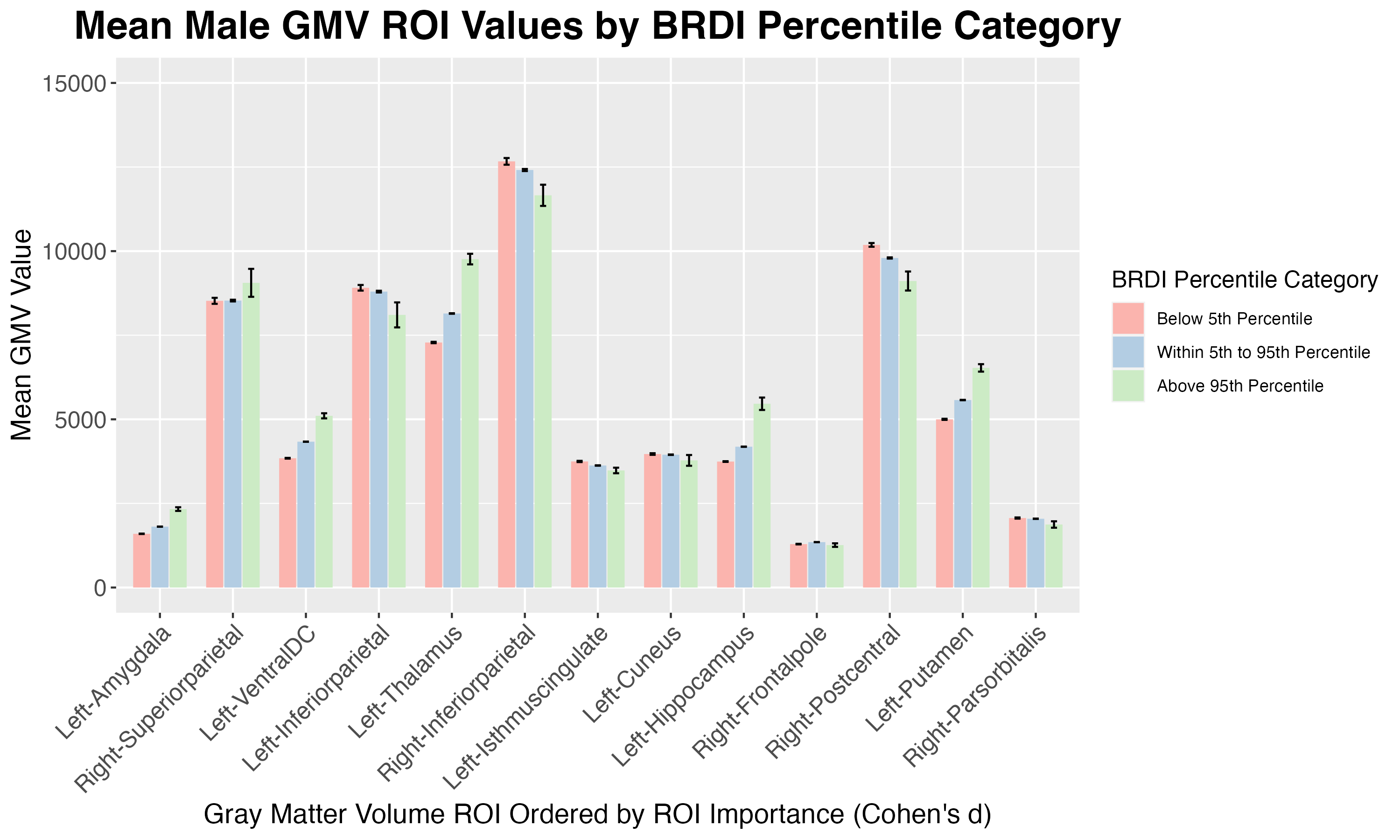

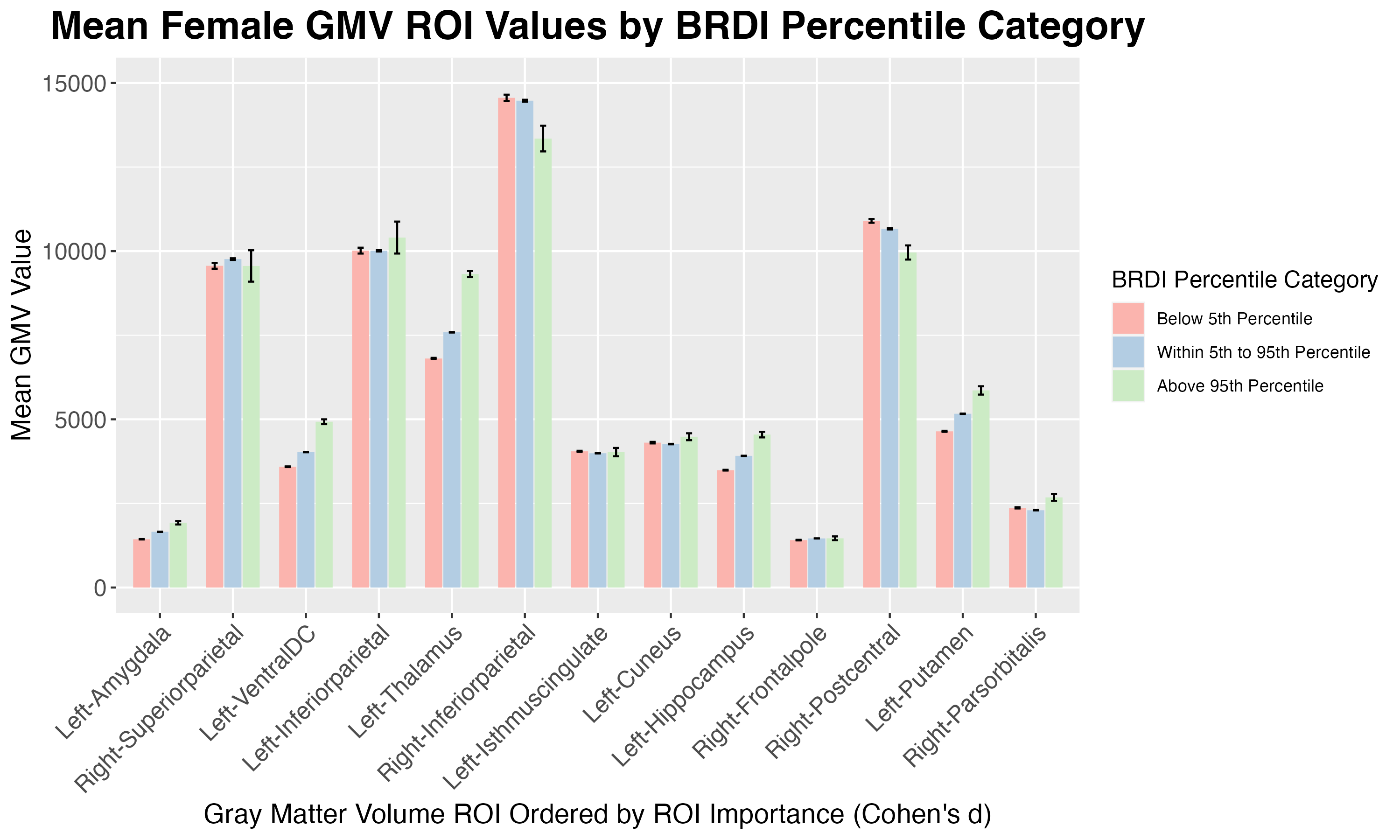

Table S2

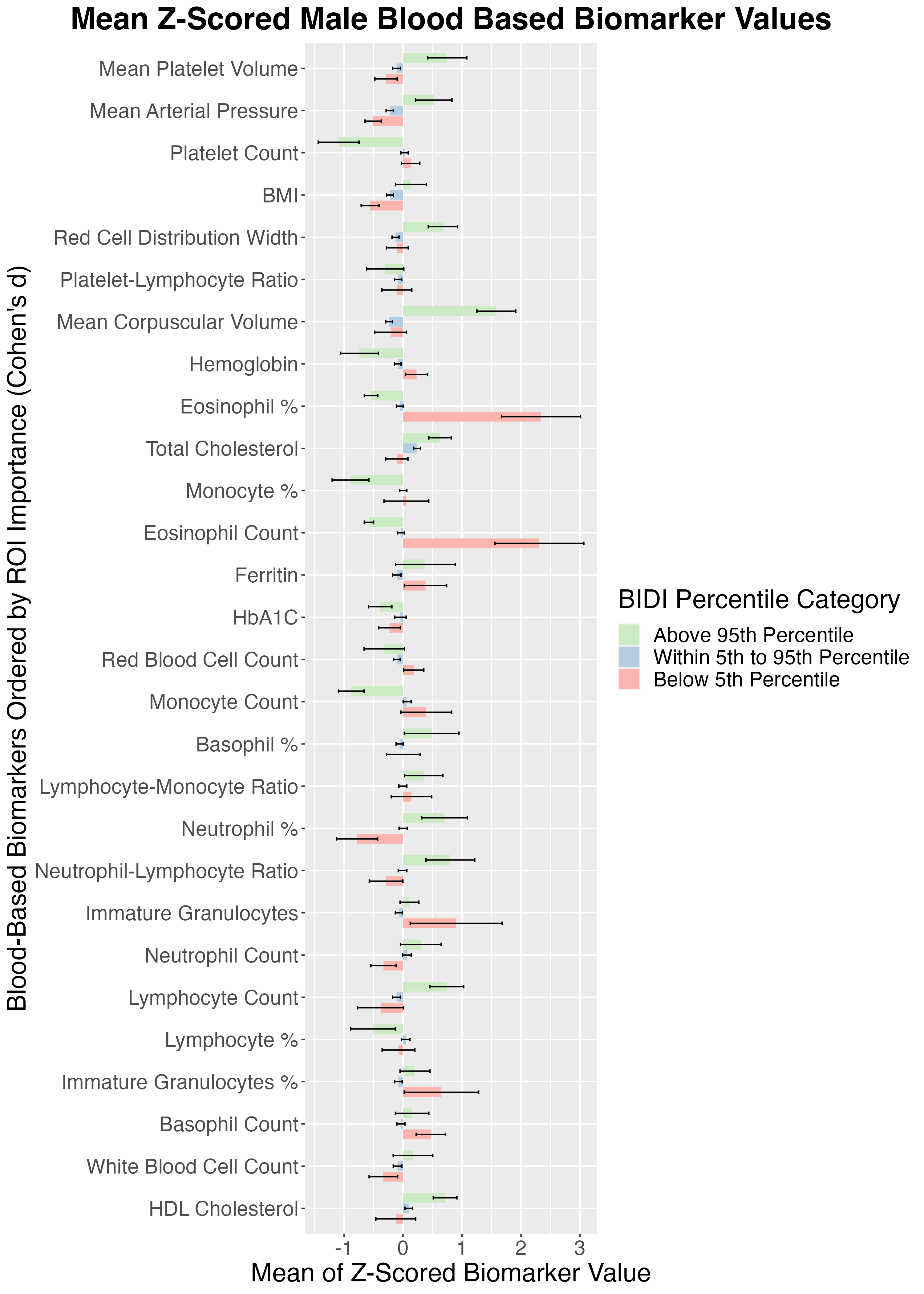

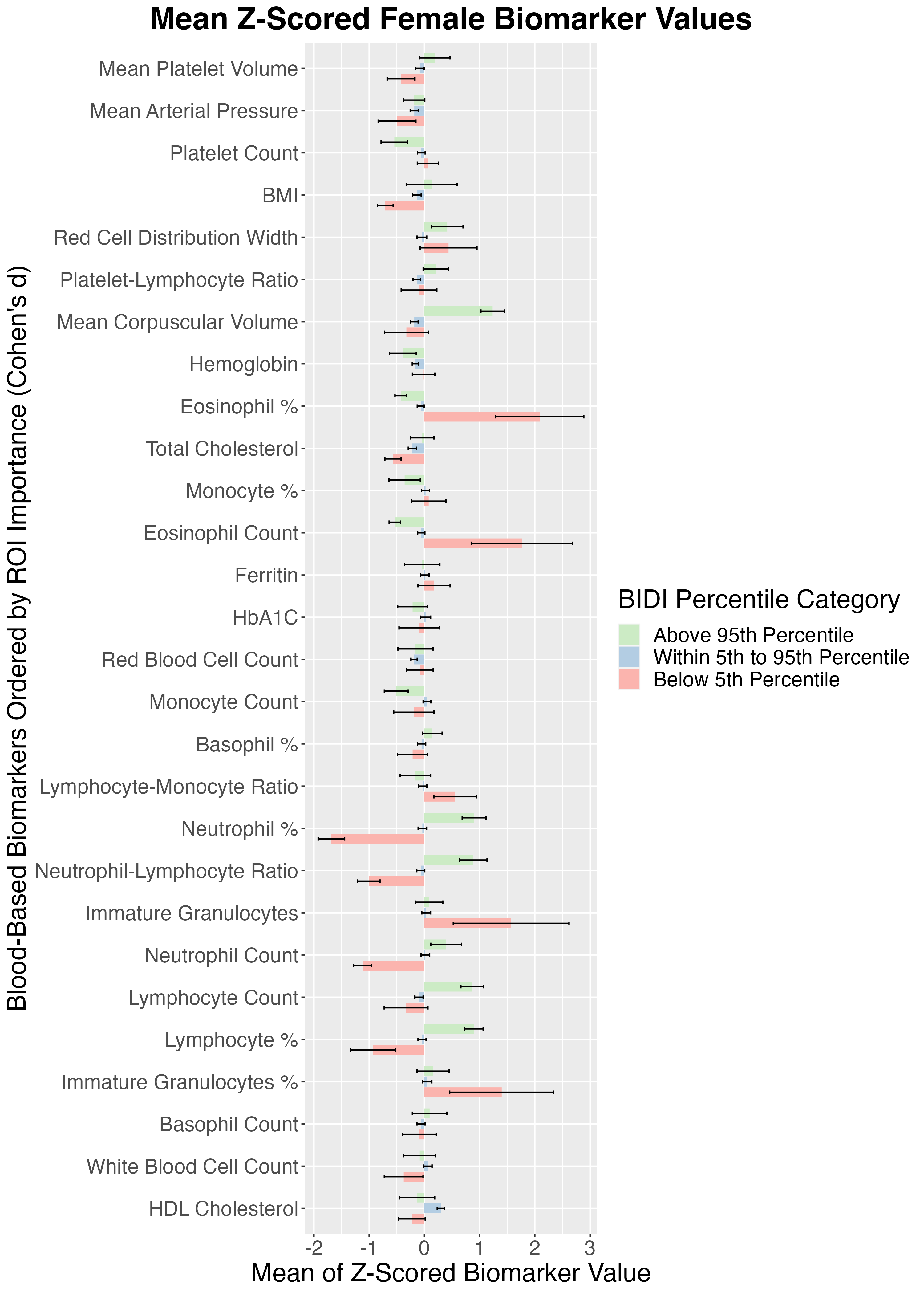

Table S3

Biological Age Year 2 (n=1846)

| Age |  | Race/Ethnicity |  | Sex |  |
| --- | --- | --- | --- | --- | --- |
| **10** | 92 | **White** | 1045 | **Male** | 972 |
| **11** | 871 | **Black** | 237 | **Female** | 874 |
| **12** | 789 | **Hispanic** | 323 |  |  |
| **13** | 93 | **Asian** | 33 |  |  |
| **14** | 0 | **Other** | 208 |  |  |
| **15** | 0 |  |  |  |  |

Biological Age Year 3 (n=430)

| Age |  | Race/Ethnicity |  | Sex |  |
| --- | --- | --- | --- | --- | --- |
| **10** | 0 | **White** | 252 | **Male** | 212 |
| **11** | 18 | **Black** | 52 | **Female** | 218 |
| **12** | 198 | **Hispanic** | 73 |  |  |
| **13** | 199 | **Asian** | 7 |  |  |
| **14** | 15 | **Other** | 46 |  |  |
| **15** | 0 |  |  |  |  |

Biological Age Year 4 (n=1075)

| Age |  | Race/Ethnicity |  | Sex |  |
| --- | --- | --- | --- | --- | --- |
| **10** | 0 | **White** | 590 | **Male** | 575 |
| **11** | 0 | **Black** | 128 | **Female** | 500 |
| **12** | 35 | **Hispanic** | 229 |  |  |
| **13** | 410 | **Asian** | 19 |  |  |
| **14** | 503 | **Other** | 109 |  |  |
| **15** | 127 |  |  |  |  |

Table S4

Brain Age Baseline (n=11563)

| Age |  | Race/Ethnicity |  | Sex* |  |
| --- | --- | --- | --- | --- | --- |
| **8** | 142 | **White** | 5986 | **Male** | 6018 |
| **9** | 5866 | **Black** | 1724 | **Female** | 5541 |
| **10** | 5434 | **Hispanic** | 2386 | **NA** | 4 |
| **11** | 121 | **Asian** | 237 |  |  |
| **12** | 0 | **Other** | 1230 |  |  |
| **13** | 0 |  |  |  |  |
| **14** | 0 |  |  |  |  |
| **15** | 0 |  |  |  |  |

Brain Age Year 2 (n=7865)

| Age |  | Race/Ethnicity |  | Sex* |  |
| --- | --- | --- | --- | --- | --- |
| **8** | 0 | **White** | 4336 | **Male** | 4206 |
| **9** | 0 | **Black** | 1059 | **Female** | 3658 |
| **10** | 353 | **Hispanic** | 1503 | **NA** | 1 |
| **11** | 3648 | **Asian** | 148 |  |  |
| **12** | 3409 | **Other** | 819 |  |  |
| **13** | 455 |  |  |  |  |
| **14** | 0 |  |  |  |  |
| **15** | 0 |  |  |  |  |

Brain Age Year 4 (n=2977)

| Age |  | Race/Ethnicity |  | Sex* |  |
| --- | --- | --- | --- | --- | --- |
| **8** | 0 | **White** | 1687 | **Male** | 1576 |
| **9** | 0 | **Black** | 316 | **Female** | 1400 |
| **10** | 0 | **Hispanic** | 600 | **NA** | 1 |
| **11** | 0 | **Asian** | 65 |  |  |
| **12** | 105 | **Other** | 309 |  |  |
| **13** | 1207 |  |  |  |  |
| **14** | 1325 |  |  |  |  |
| **15** | 340 |  |  |  |  |
